# Heart Snapshot: a broadly validated smartphone measure of VO_2_max for collection of real world data

**DOI:** 10.1101/2020.07.02.185314

**Authors:** Dan E. Webster, Meghasyam Tummalacherla, Michael Higgins, David Wing, Euan Ashley, Valerie E. Kelly, Michael V. McConnell, Evan D. Muse, Jeff Olgin, Lara Mangravite, Job Godino, Michael Kellen, Larsson Omberg

## Abstract

Expanding access to precision medicine will increasingly require that patient biometrics can be measured in remote care settings. VO_2_max, the maximum volume of oxygen usable during intense exercise, is one of the most predictive biometric risk factors for cardiovascular disease, frailty, and overall mortality.^1,2^ However, VO_2_max measurements are rarely performed in clinical care or large-scale epidemiologic studies due to the high cost, participant burden, and need for specialized laboratory equipment and staff.^3,4^ To overcome these barriers, we developed two smartphone sensor-based protocols for estimating VO_2_max: a generalization of a 12-minute run test (12-MRT) and a submaximal 3-minute step test (3-MST). In laboratory settings, Lins concordance for these two tests relative to gold standard VO_2_max testing was *p*_c_=0.66 for 12-MRT and *p*_c_=0.61 for 3-MST. Relative to “silver standards”^5^ (Cooper/Tecumseh protocols), concordance was *p*_c_=0.96 and *p*_c_=0.94, respectively. However, in remote settings, 12-MRT was significantly less concordant with gold standard (*p*_c_=0.25) compared to 3-MST (*p*_c_=0.61), though both had high test-retest reliability (ICC=0.88 and 0.86, respectively). These results demonstrate the importance of real-world evidence for validation of digital health measurements. In order to validate 3-MST in a broadly representative population in accordance with the *All of Us* Research Program^6^ for which this measurement was developed, the camera-based heart rate measurement was investigated for potential bias. No systematic measurement error was observed that corresponded to skin pigmentation level, operating system, or cost of the phone used. The smartphone-based 3-MST protocol, here termed *Heart Snapshot*, maintained fidelity across demographic variation in age and sex, across diverse skin pigmentation, and between iOS and Android implementations of various smartphone models. The source code for these smartphone measurements, along with the data used to validate them,^6^ are openly available to the research community.

## Background

Clinical decisions and public health policy are increasingly informed by multifactorial analysis of large cohorts of patients and their associated outcomes. Traditionally, cardiovascular health has been assessed using risk scores such as the Framingham Risk Score^7^, Reynolds Risk Score^8^, Qrisk^9^ and others that integrate multiple factors including demographic data, comorbidities, and biometrics paired to imaging-based assessments measuring vascular blockage and blood flow in higher-risk and symptomatic individuals. While these factors have clear correlation to cardiovascular health, their inclusion in integrative risk calculations was promoted in part because they can be rapidly evaluated across many individuals. However, one of the most predictive biometrics for cardiovascular health^10^ and overall mortality^1^, VO_2_max, is typically not incorporated in these risk calculators due to the burdens associated with obtaining this measurement.^3,4^

Cardiorespiratory fitness as measured by VO_2_max represents the integrated function of physiological systems involved in transporting oxygen from the atmosphere to the skeletal muscles to perform physical work. Existing gold standard techniques for measuring VO_2_max are based on protocols that use exercise on a treadmill or stationary bicycle paired with direct measurement of oxygen consumption at various workloads including maximal exertion.^11,12^ However, the requirement to exercise at the maximal aerobic threshold limits deployment in some populations for safety reasons, and the need for specialized equipment and personnel has prohibited widespread adoption of VO_2_max testing in research and clinical settings.

Because of these limitations of gold standard VO_2_max measurements, numerous “silver standard”^5^ VO_2_max estimation protocols have been developed that rely on simpler equipment or submaximal levels of exertion. These protocols trade off measurement accuracy for ease of deployment in a wider range of settings and for populations with differing levels of capacity.^13^ However, these protocols were typically developed and validated in small, homogeneous populations and some subsequent validation studies have been criticized for demonstrating participant selection bias.^14^ To overcome these limitations, we sought to develop a digital VO_2_max estimation protocol that could be self-administered remotely using only the sensors within a smartphone, and we sought to validate this measure within a broadly representative population.

## Results

Two silver standard VO_2_max estimation protocols were chosen as a basis for developing the smartphone tests. The first is the Cooper protocol,^15^ consisting of a 12-minute walk/run test (12-MRT) where individuals cover as much distance as possible in 12 minutes on a flat course. The Cooper protocol estimates VO_2_max from the total distance traveled during the 12 minutes. The other is the Tecumseh protocol,^15,16^ which consists of a 3-minute submaximal step test (3-MST) where individuals step up and down an 8-inch step at a constant rate for 3 minutes. In the 3-MST protocol, VO_2_max is estimated from heart rate measurements during the recovery period. In adopting these protocols for smartphones, we developed self-guided instructions (Supplemental figure 4) with the GPS to record distance during 12-MRT, and the smartphone camera to record heart rate during recovery for the 3-MST.

To assess the validity of these approaches, gold standard VO_2_max treadmill testing was performed with 101 participants distributed across age deciles 20-80 (Table 1). Every participant also performed the silver standard and smartphone 12-MRT and 3-MST protocols in the clinic, with 3 instances of each smartphone protocol performed in a remote setting (Figure 1a).

**Table 1.** Demographic landscape and gold standard measures of maximum heart rate and VO_2_max in the validation cohort.

| Age Decile | Sex<br>*N* (Female\|Male) | Age<br>(mean +/- std) | Ethnicity<br>(% White) | BMI<br>(mean +/- std) | Max HR<br>(bpm) | VO2max<br>(mL/kg/min) |
| --- | --- | --- | --- | --- | --- | --- |
| (20, 30] | 16 (7\|9) | 24.6 +/- 3.2 | 43.8 | 24.0 +/- 2.9 | (193) | (47) |
| (30, 40] | 20 (14\|6) | 35.5 +/- 3.1 | 65.0 | 24.0 +/- 3.5 | (186) | (44) |
| (40, 50] | 20 (13\|7) | 44.7 +/- 3.0 | 65.0 | 26.8 +/- 5.4 | (181) | (39) |
| (50, 60] | 19 (8\|11) | 55.3 +/- 2.9 | 68.4 | 27.0 +/- 3.9 | (174) | (40) |
| (60, 70] | 14 (11\|3) | 64.8 +/- 3.2 | 85.7 | 28.5 +/- 4.7 | (160) | (31) |
| (70, 80] | 5 (3\|2) | 75.0 +/- 3.4 | 100.0 | 27.1 +/- 3.0 | (141) | (31) |

**Figure 1.**
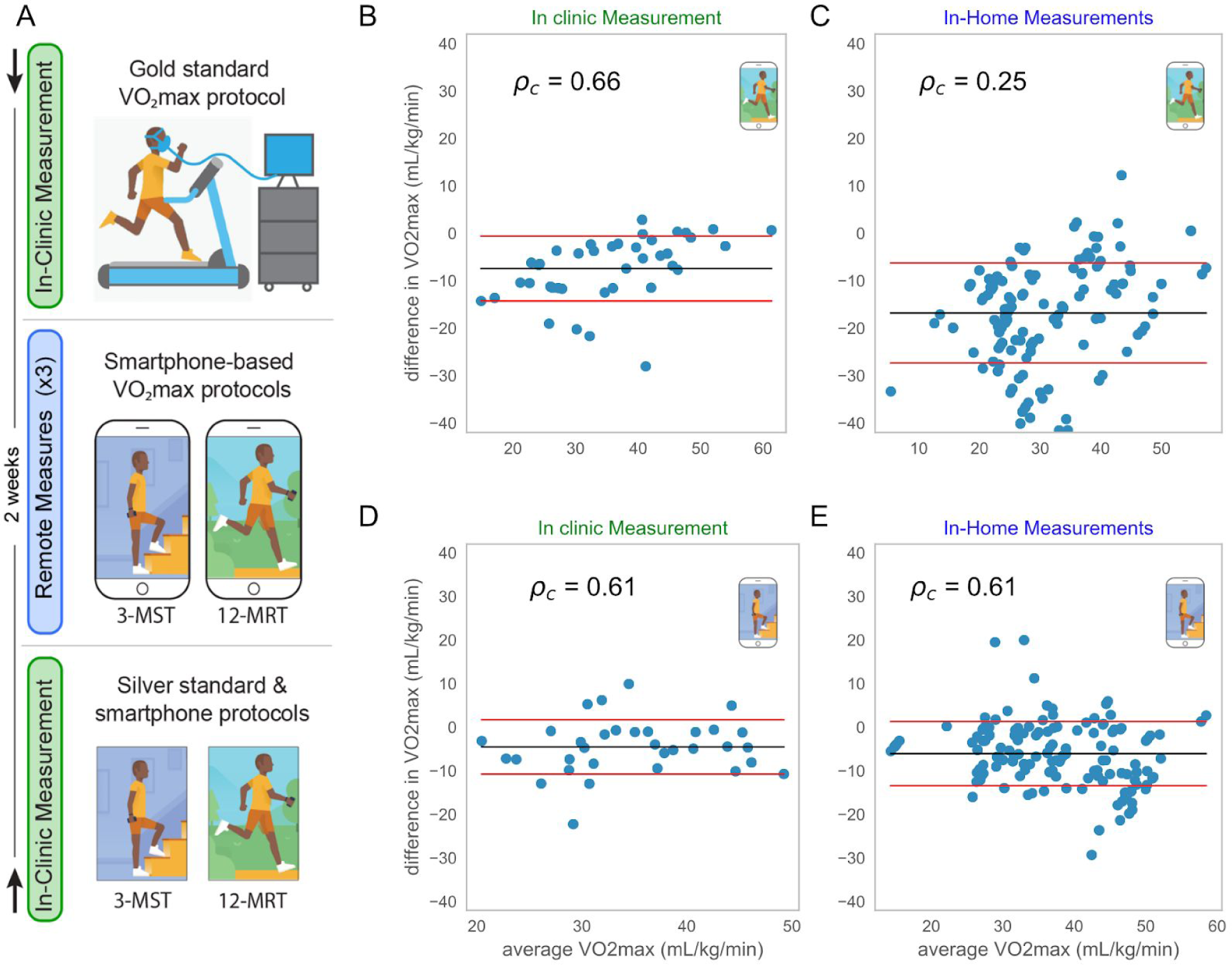
Validation protocol and primary results of validation. (A) Participants in the study were randomized into two groups. The first group (denoted by the downward-facing arrow at top) performed a gold standard VO_2_max protocol and received training on day 1. The second group performed the two silver standard protocols concurrently with the smartphone protocols on day 1 (denoted by the upward-facing arrow at bottom). Both groups then performed the two smartphone protocols remotely up to three times during a two week period. B-E show Bland-Altman plots comparing the gold standard VO_2_max with smartphone measures from: B) 12-MRT performed in clinic, C) 12-MRT performed remotely (up to 3 repeats per participant), D) 3-MST in clinic, and E) 3-MST remotely. For upper limits of performance comparing non-smartphone measures to gold standard see supplemental figure 1.

In-clinic 12-MRT distance was measured on a 400m track and by the smartphone GPS. In-clinic heart rate was measured via radial pulse measured by trained research staff, a chest-worn Polar heart monitor, a wrist-worn Fitbit Charge 2, and the smartphone camera with the flash activated. Comparisons between gold standard, silver standard, and smartphone-based protocols for VO_2_max estimation were performed using Bland-Altman analysis^17^ and Lin’s concordance index (*p*_c_). The concordance between gold standard VO_2_max and the silver standard Cooper protocol (*p*_c_=0.61, Supplemental Figure 1a) and the silver standard Tecumseh protocol (*p*_c_=0.70, Supplemental Figure 1b) were in line with previously published results.^18,19,20^ Concordance of smartphone-based protocols with gold standard VO_2_max testing was *p*_c_=0.66 for 12-MRT (Figure 1b) and *p*_c_=0.61 for the 3-MST (Figure 1d). Concordance of smartphone-based protocols with silver standard protocols was *p*_c_=0.96 for 12-MRT and *p*_c_=0.94 for the 3-MST. These results demonstrate that the smartphone-based protocols fall short of recapitulating gold-standard VO_2_max testing, but are highly concordant with validated silver standard VO_2_max estimation protocols in a laboratory setting.

To investigate whether the concordance of in-clinic measurements would generalize to remote and unsupervised settings, the smartphone protocols were also performed up to 3 times at home by each participant. We observed an approximately equal test-retest reliability between the two tests (3-MST ICC=0.86, 12-MRT ICC=0.88). However, while the 3-MST translated well to an unsupervised setting (*p*_c_=0.61, Figure 1e), the 12-MRT demonstrated a pronounced drop in concordance (*p*_c_=0.25, Figure 1c), despite a highly accurate distance measurement from the smartphone (*p*_c_=0.96) based on comparisons made in a clinical setting.

As the 12-MRT is dependent on maximal effort, participants were surveyed directly after their run about their performance. In 137 of 216 runs performed remotely (63.4%), participants reported the run to be “their best effort”. Therefore, only these 137 runs were used to estimate VO_2_max in our analysis. Supplemental figure 1 captures the results of all 216 runs subdivided by self-reported effort. While the context-dependent failure of the 12-MRT in remote settings may be attributable to many factors, this result highlights the importance of both clinical and unsupervised real-world evidence for the validation of novel digital health measurement modalities.

The smartphone-based 3-MST protocol, hereafter referred to as *Heart Snapshot*, was generalizable between clinical and remote assessments, and was robust over a large range of fitness levels. Being robust to unsupervised measurements is necessary to achieve the scale intended for the targeted 1 million participants in *All of Us* Research Program (AoURP)^6^, which will employ a ‘bring your own device’ strategy for remote self-measurement of VO_2_max. AoURP also aims to recruit a study population matching the full demographic diversity of the US, emphasizing inclusion of groups often underrepresented in biomedical research, such as ethnic and racial minorities. As prior studies have shown differing results as to whether optical techniques for heart rate detection (photoplethysmography) can be demographically biased, ^21,22^ we sought to investigate any differences in *Heart Snapshot* accuracy across variation in skin tone. A followup calibration study for heart rate measurements was conducted with 120 participants distributed approximately evenly across defined Fitzpatrick skin types,^23^ using 8 different smartphones (3 iPhone, 5 Android ranging in cost from $99 to $999 at the time of writing). These phones were chosen to be representative of different operating systems, quality of sensors, speed of processing, and camera configurations. Importantly, we observed no significant difference in heart rate measurement accuracy between categorical Fitzpatrick skin types or systematic measurement error proportional to skin color at either end of the spectrum (Figure 2A).

**Figure 2.**
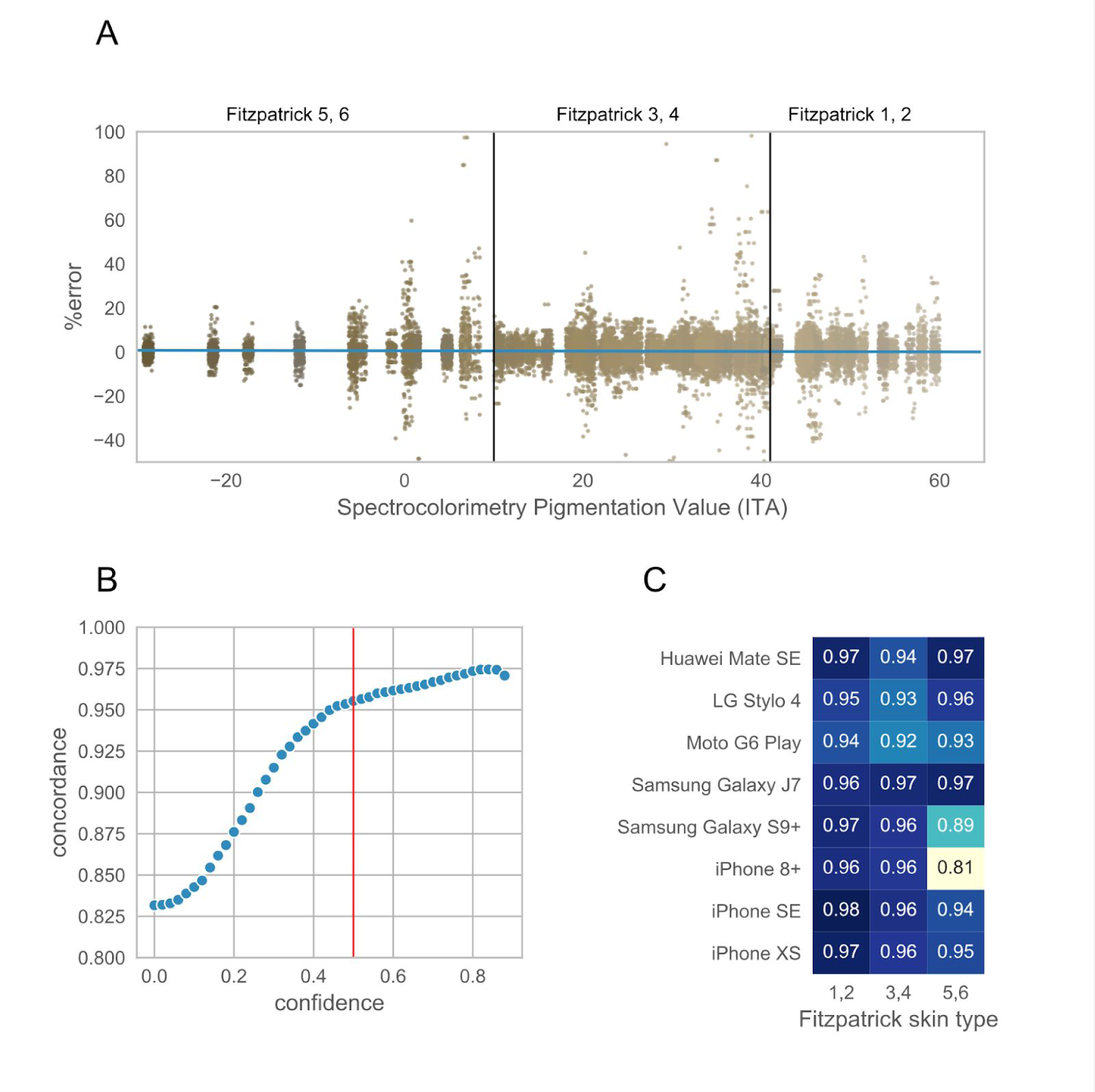
Validation of heart rate measurements across different skin tones and hardware configurations in the calibration study. A) Percent error in heart rate estimation from ground truth as a function of different colors captured by spectrocolorimetry under the jaw. Each dot represents a 10 second window of heart rate in one individual. B) Distribution of concordance between heart rate using pulse oximetry and smartphone as the confidence cutoff is changed. Red line represents the chosen cutoff used for analysis (supplementary figure 2). C) Concordance as a function of smartphone models and Fitzpatrick skin tones.

To facilitate quality control of the measurements across different smartphones, a confidence score was developed to provide a readout of the quality of the heart rate measurements. This confidence score is derived from the autocorrelation function (ACF) of heart rate signal across 10-second measurement windows. Using the calibration study results, a balance between quality of measurements was weighed against loss of data by choosing a filtration cutoff at a confidence >=0.5. This resulted in a *p*_c_=0.95 in the calibration cohort between a pulse-oximetry pulse measurement and the camera-estimated heart rate (Figure 2b). In selecting this confidence score as a cutoff, we observed that 81.4% of all measurement windows were retained in this calibration cohort (Supplementary Figure 2a). This same cutoff was used in the above validation of *Heart Snapshot* where the heart rate concordance with a chest-worn Polar heart monitor was *p*_c_=0.95 and *p*_c_=0.83 when compared to a wrist-worn Fitbit Charge 2 both at home and in the clinic. This can be compared to *p*_c_=0.92 between the Polar and Fitbit Charge 2 (Supplemental Figure 3).

Taken together, *Heart Snapshot* heart rate measurements in any of the combinations of the Fitpatrick skin tones and 8 smartphones used in the calibration study resulted in a concordance greater than or equal to *p*_c_=0.81 (Figure 2c), which is in line with previous smartphone-based modalities for heart rate monitoring.^24^ Importantly, performance did not correlate with device cost, with all phones selling for under $200 dollars performing better than *p*_c_>0.92 for any skin tone.

In summary, *Heart Snapshot* measured VO_2_max with similar accuracy to supervised, in-clinic tests such as the Tecumseh or Cooper protocols, while also generalizing to remote and unsupervised measurements. *Heart Snapshot* measurements demonstrated fidelity across demographic variation in age and sex, across diverse skin pigmentation, and between various iOS and Android phone configurations. This software is freely available (see methods) with all validation data and analysis code (https://github.com/Sage-Bionetworks/CRF_validation_analysis)

## Discussion

While multiple devices can estimate VO_2_max, including several currently marketed consumer devices,^25^ the underlying data and algorithms are usually not published. The lack of data and method transparency limits the utility of these approaches for discovery-based research where reproducibility is paramount. In contrast, an open approach to method validation can also serve as a foundation for downstream research in different conditions or populations to generate normative data for interpreting results.^26^

As many dedicated hardware devices for digital health in the consumer sphere have experienced short half-lives of availability, we believe that the dependency only on a smartphone with a flash and camera may provide a greater degree of ‘future-proofing’ for *Heart Snapshot*. This will be important for consistent, longitudinal measurements that may uncover patterns of VO_2_max variance over time, especially in very large scale studies such as the AoURP.

*Heart Snapshot* attempts to maximize concordance with gold standard methods for estimating VO_2_max, but it is worth noting that this analysis used an existing validated algorithm^18^ that was based on in-clinic procedures and measurement tools. *Heart Snapshot* could become more personalized than traditional protocols, for example adapting to a participant’s maximum step cadence as measured by smartphone accelerometry. Further concordance with gold standard measures may be achieved by optimizing the parameters of the traditional algorithm or including new variables, but this will require a distinct cohort for testing any models that have been trained on this dataset.

The emerging development of consumer technology provides us with unprecedented opportunities to evaluate the utility of additional digital biomarkers to improve risk management strategies for population health and for precision health at the level of an individual. Paired with access to large population studies, such as the *All of Us* Research Program^6^ that collect health questionnaires, electronic health records, physical measurements, biospecimens, and digital health technology data, we can rapidly test emerging digital health measures for their potential to advance precision medicine.

## Acknowledgements

We would like to acknowledge Steve Steinhubl, Shannon Young, Nathaniel Brown, Joshua Liu, Erin Mounts, Stockard Simon, and Woody MacDuffie for their contributions to this work.

## Author Contributions

DEW and LO wrote the first draft of the paper. LO, MK and JG developed the study and protocol, MT developed the algorithms for heart rate measurements LO and MT performed the analysis. MH, DW and JG recruited the participants and performed all measurements in the lab. MK and DEW oversaw the design and development of the heart snapshot applications. EA, VEK, MVM, EDM, JO and LM helped identify the protocols for generalization, provided expert input and edited the paper together with MK, LO, JG, DW, MT, MH and DW. LO and MK contributed equally to this effort.

## Supplementary Data

- Supplemental **Table S1** complete EPARC metadata

**Supplemental Figure 1.**
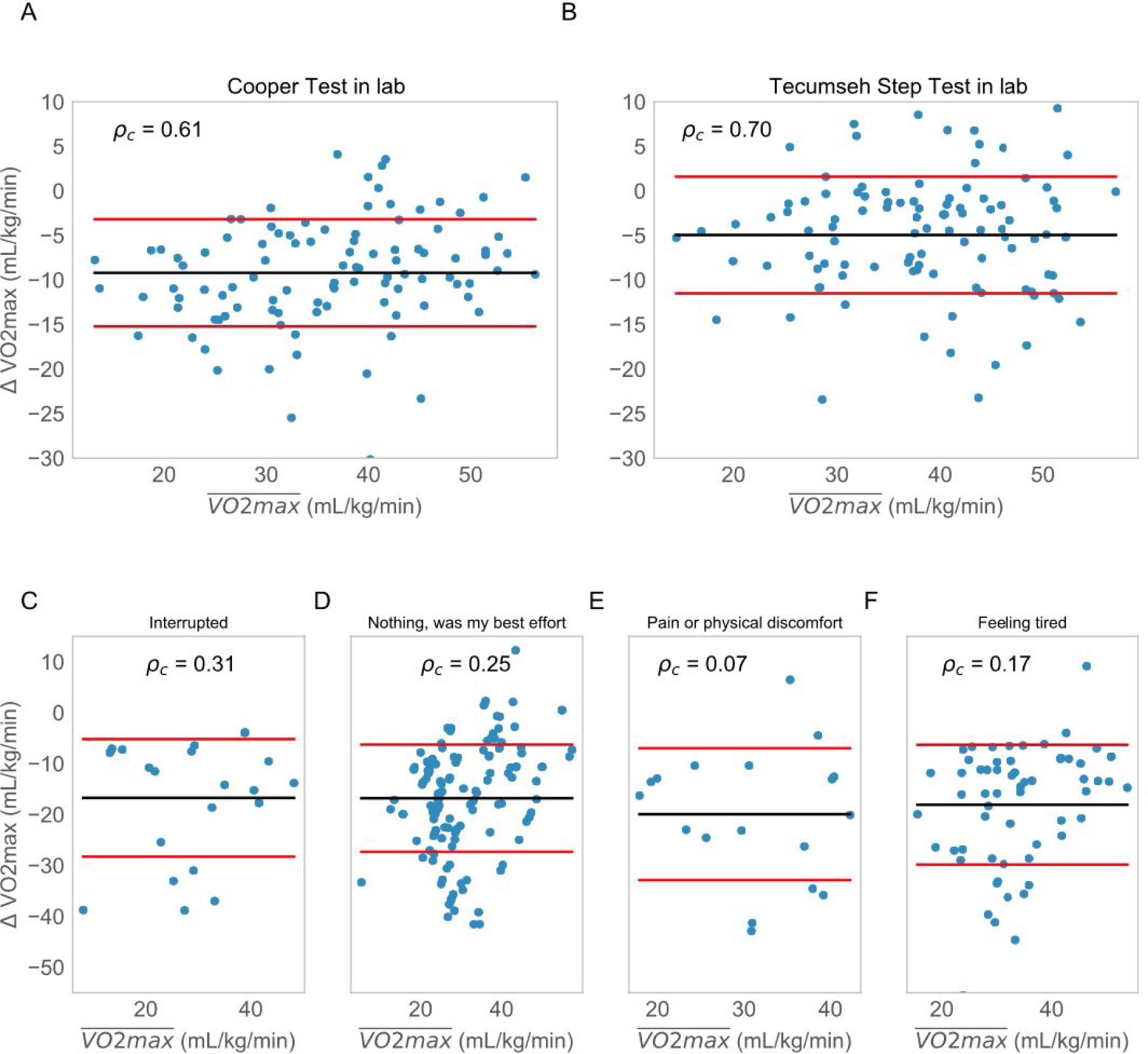
Comparison of in-clinic performance of silver standard protocols relative to gold standard for A) 12-MRT and B) 3-MST. For each plot we are showing the difference between the ground truth VO_2_max measurement and measurements obtained using the distance run around a track (for A) and heart rate via radial pulse measured by trained research staff (for B). This distance was also measured using GPS and heart rate was measured using a chest strap and Fitbit. The concordance between distance measured around the track and measured using the GPS in the phone was 0.96. C-F) shows the concordance of the 12-MRT test for different values of self-reported effort.

**Supplemental Figure 2.**
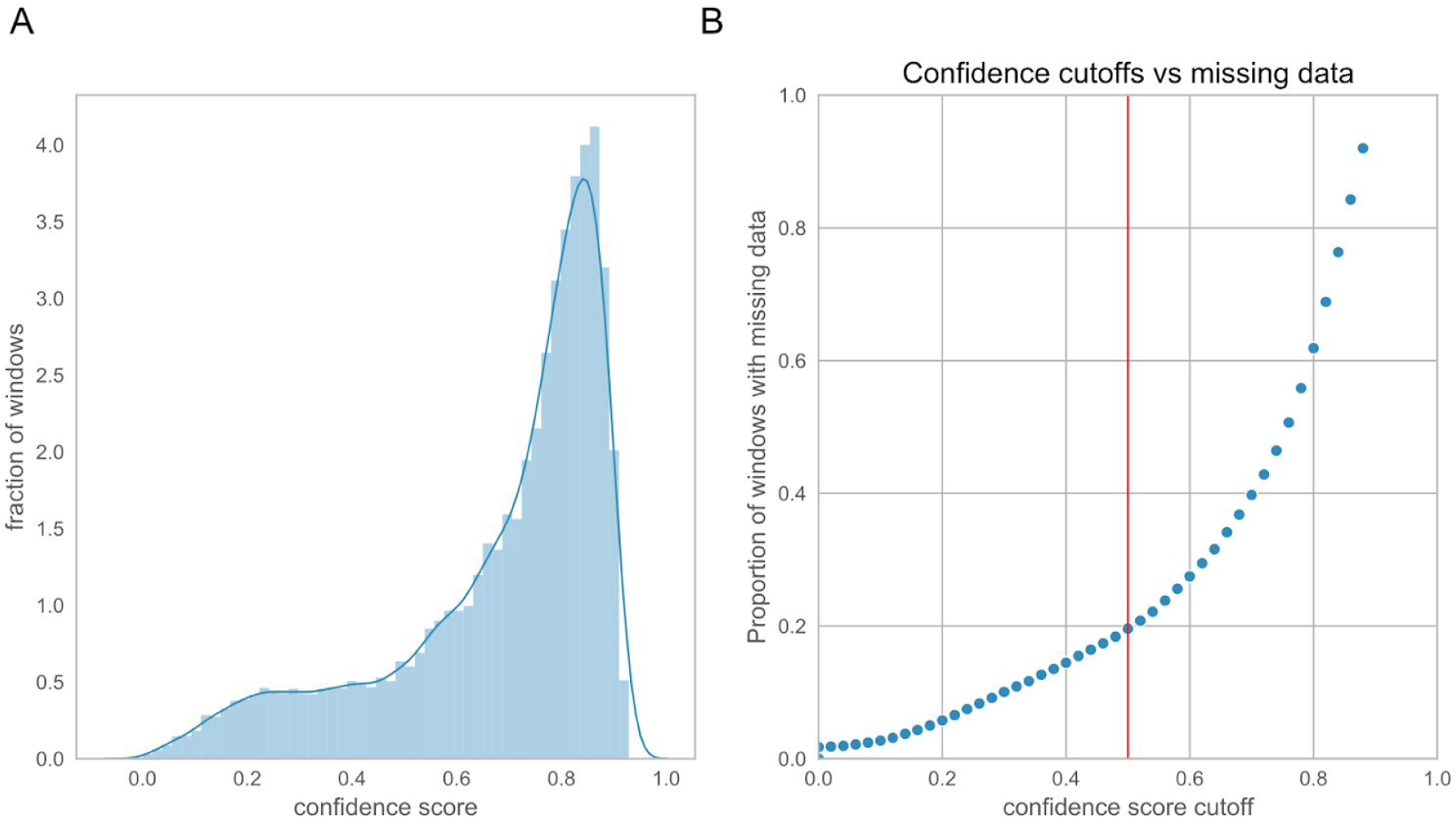
Effect of different confidence cutoff on the amount of missing data from the calibration study. A) Distribution of best confidence across red and green channels in the calibrations study and B) percent of the 10 second windows that are filtered out at different cutoffs of the confidence score. The cutoff used in the analysis is 0.5 marked by the red line.

**Supplemental Figure 3.**
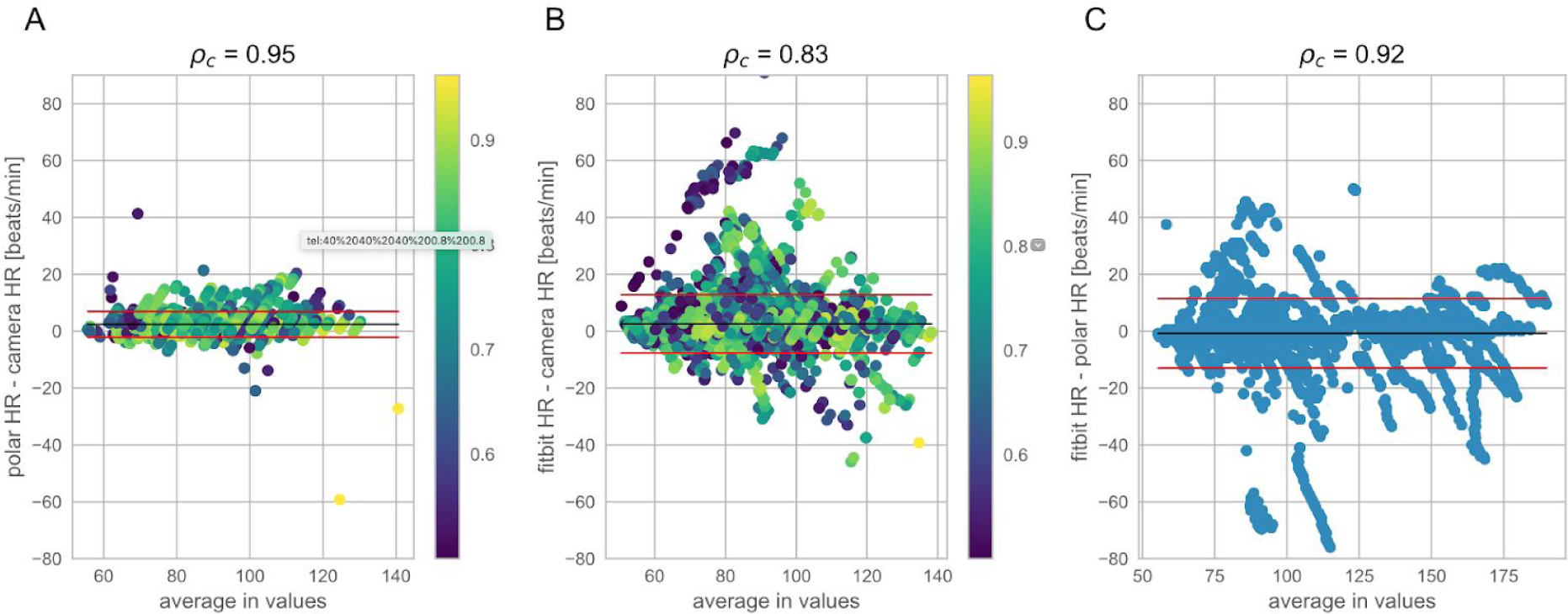
Bland-Altman analysis comparing heart rate measurements in the *validation study* using data collected during the Tecumseh tests. In the validation cohort, participants used multiple ways of collecting heart rate. The method being tested, the smartphone camera, was compared to: A) a Polar chest strap while in the clinic when both were used and B) a Fitbit worn during the entirety of the study. C) We also compared the Polar strap to the Fitbit for all time that both were worn.

**Supplemental Figure 4.**
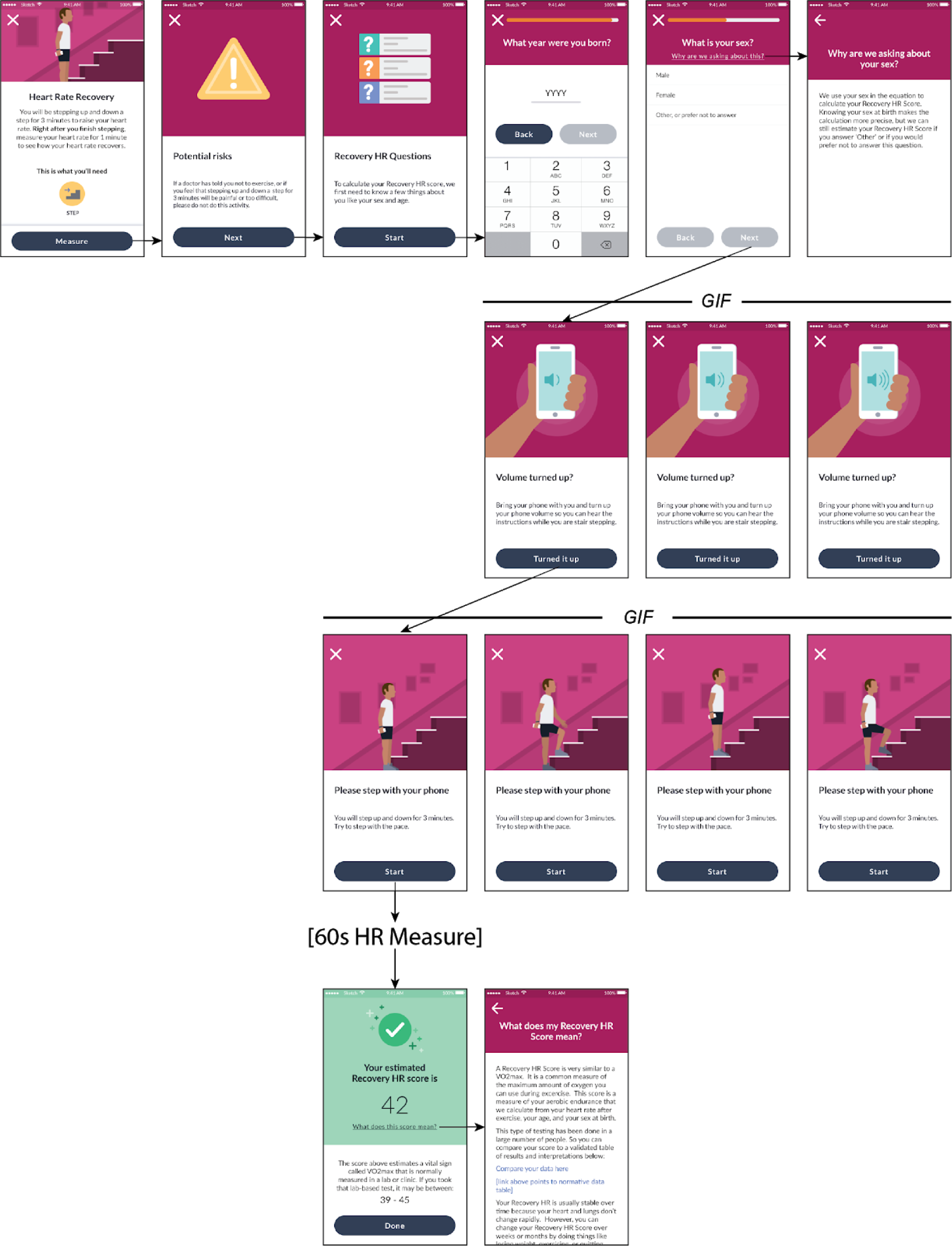
Self-guided instructions and screen workflow for performing the Heart Snapshot VO_2_max estimate.

## Methods

### VO_2_max Validation Cohort Procedures and Measures

All study procedures were approved by the University of California, San Diego (UCSD) Institutional Review Board (approval number 171815). All participants provided written informed consent and attended two in-person study visits at the Exercise and Physical Activity Resource Center (EPARC).

A convenience sample of 101 adults between 20 and 79 years of age were recruited, largely balanced across age deciles and sex (Supplementary Table S1). Potential participants were contacted by trained EPARC staff via email or telephone, and they underwent a screening to ascertain their eligibility. Participants were included if they were 1) able to consent and participate in the study using English; 2) between 20 and 79 years of age; 3) willing and able to attend two in-person study visits that included either a VO_2_max test or a 12-MRT and a 3-MST within a two-week period; 4)willing and able to undertake up to three 12-MRT and 3-MST at home over a two-week period; 5) willing and able to download the smartphone application developed to measure cardiorespiratory fitness on a compatible Android or iOS device and use it during all tests over a two-week period; and 6) willing and able to download Fitbit’s smartphone application on a compatible Android or iOS device and connect and wear a study-provided Fitbit Charge 2 during all tests over a two-week period. Participants were excluded if they 1) were greater than 12 weeks pregnant; 2) had a heart or cardiovascular condition, including coronary artery disease, congestive heart failure, diagnosed abnormality of heart rhythm, atrial fibrillation, and/or a history of myocardial infarction; 3) required use of an external device to assist heart rhythm (e.g., a pacemaker); 4) had a serious respiratory disease, including chronic obstructive pulmonary disease, exercise induced asthma, and/or pulmonary high blood pressure; 5) required use of supplemental oxygen; 6) required use of a beta blocker or other medications known to alter heart rate; and 7) answered “yes” to one or more questions in the American College of Sports Medicine’s Physical Activity Readiness Questionnaire (PAR-Q) and/or report two or more risk factors for exercise testing and did not receive subsequent medical clearance. The PAR-Q is a widely-accepted tool used to assess an individual’s fitness for tests involving cardiovascular exercise^27^.

Upon completion of the telephone screening (and if necessary, receipt of medical clearance), potential participants were scheduled to attend the first testing session at UCSD. They were asked to report to the testing session well-hydrated and in athletic attire. Participants were guided through the process of downloading and installing the smartphone application developed to measure cardiorespiratory fitness, as well as Fitbit’s smartphone application, and they were fitted with a wrist-worn Fitbit Charge 2 according to the manufacturer’s recommendations. Participants were asked to provide their age, sex at birth, ethnicity, and race. Weight (to the nearest 0.1kg) and height (to the nearest 0.1cm) was measured using a calibrated digital scale and stadiometer (Seca 703, Seca GmbH & Co. KG., Hamberg, DE). Both weight and height were measured with participants wearing lightweight clothes but without shoes, and two separate measurements were averaged (if weight or height measurements differed by more than 1%, then a third measure was taken and the average of the two measures that differed by less than 0.2kg or 0.5cm, respectively, were used).

At the first testing session, participants either undertook 1) a VO_2_max test or 2) an in-clinic 3-MST and 12-MRT. A randomization procedure implemented prior to the scheduling of the first testing session determined which test procedure participants undertook at the first testing session. Participants were then expected to complete the other test procedure during the second testing session.

### Treadmill-based, gold standard VO_2_max measurement

Participants completed a maximal graded exercise test on a Woodway 4Front treadmill (Woodway, Waukesha, WI) that was calibrated monthly for accuracy of speed and grade. The maximal graded exercise test protocol began with a warm-up at a self-selected pace on the treadmill for 5 to 10 min. During the warm-up, EPARC staff explained how to use the Borg Rating of Perceived Exertion (RPE) scale and reminded participants that they were expected to achieve their maximal level of exertion.^28^

Participants were then equipped with a breath mask that covers the nose and mouth (KORR Medical Technologies, Salt Lake City, UT), and a Bluetooth enabled heart rate monitor worn on the chest (Garmin, Olathe, KS). The preprogrammed treadmill protocol began with participants running at 5 mph with 0% incline for 3 min. The workload was then increased approximately 0.75 Metabolic Equivalent of Tasks (METs) every minute. This was achieved via an increase in speed (0.5 mph/min) each minute until the participant was 0.5 mph above their self-determined comfortable speed or until a maximal speed of 9.0 mph was reached. If the participant’s capacity allowed them to continue beyond this upper speed limit before reaching volitional fatigue, then the treadmill speed was kept constant, but the grade (i.e., incline) of the treadmill was increased by 1.0% each minute until volitional fatigue was reached. RPE was assessed during the final 10 s of each minute, and the protocol continued until the participant signaled to stop (i.e., indication of volitional fatigue). Upon indication of volitional fatigue, the treadmill was immediately slowed to 2.0 mph, and participants were encouraged to walk until completely recovered. Breath by breath oxygen uptake (VO_2_) was continuously measured using an indirect calorimeter (COSMED, Trentino, Italy) that was calibrated for gas volume and fractional composition immediately (i.e., less than 30 min) before the start of the maximal graded exercise test protocol.

### Tecumseh Test (3-MST) and Cooper Test (12-MRT) In-clinic procedure

All participants were fitted with a chest-worn heart rate monitor (Polar, Finland) that was used for real-time monitoring by trained EPARC staff throughout both the 12-MRT and 3-MST. For the 3-MST, participants were instructed to step up and down from a single step 8 inches in height at a rate of 24 steps per minute for 3 minutes.^29^ The cadence of stepping was monitored by trained EPARC staff. Radial pulse was measured from second 31 to second 60 after the 3 minutes of stepping. Upon completion of the test, participants were asked to sit in a chair and rest. After a minimum of 10 minutes of rest, participants then completed a 5-minute self-determined light intensity warm-up. They were then instructed to cover as much distance as possible on a flat 400m track for 12 minutes. The distance traveled was measured after the 12 minutes.^15^

### Distance estimation using privacy-preserving GPS data

The distance recorded by the smartphone during the 12-MRT was validated against the actual distance. The smartphone recorded displacement information sampled at 1Hz which consists of relative location measurements, i.e. the change in location with respect to the last recorded measurement. The iPhones measured displacement in meters whereas on Android measured relative changes in latitude and longitude — requiring an estimate of the absolute latitude and longitude to be added back into the measurements to obtain an accurate estimate of distance.

The first distance estimation method entailed summing the euclidean distances between subsequent GPS points. Since GPS measurements have a range error dependent on atmospheric effects and numerical errors, a second method was employed which computed distance after smoothing the trajectory of the GPS path using a Savitzky-Golay smoothing filter.

### Camera-Based Heart Rate Estimation

Blood flow through the fingertip was measured through video with the rear-facing camera while the flash was on. Resting heart rate was captured with 20 seconds of recording while the 3-MST required 60 seconds of recording. During the capture we found it was important to fix the focal length to infinity, turn off any HDR (if applicable), and set the frame rate (fs) to 60 Hz if possible and if not the default highest allowed by the phone. To preserve privacy we didn’t record the video but instead summarized each video frame to the mean of all pixel intensities per color channel in RGB space.

These intensities yielded three times series, one for each color. These time series were filtered and mean centered before being split into shorter 10 second windows. By assuming a periodic signal for these windows the autocorrelation function (ACF) was used to estimate the period by finding the peaks and their corresponding lags. The relative magnitude of the peaks to the maxima of the ACF was used to generate a confidence score, which quantifies the extent to which the signal is periodic or if the peak at the fundamental frequency (i.e the peak with the highest magnitude) is a spurious peak. The ACF is calculated over a 10s window, as this provides sufficient heart beat observations post processing to estimate heart rates ranging from 45-210BPM.

To filter potentially spurious peaks, a magnitude threshold relative to the magnitude of the peak at zero-lag was employed. The confidence score was calculated as the ratio of the magnitude of the peak corresponding to the fundamental frequency to the next peak. The confidence score is roughly an indicator of how periodic the signal is, a property indicative of the heart rate signal in a short finite time window. The different color channels were merged by choosing the heart rate estimate from the channel (red or green) that has the maximum confidence score within a given window.

### Estimation of VO_2_max

#### 3-MST

Multiple formulae for predicting VO_2_max from the Tecumseh step test and its variations have been developed,^16,30^ here we use the following established by Milligan^30^ 

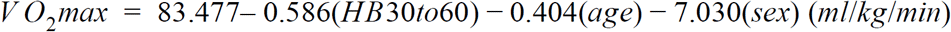

Where, *HB*30*to*60 is the number of beats between *30* to *60* seconds post step test, *age* is the age of the subject, and *sex* is *0* if male and *1* if female.

#### 12-MRT

VO_2_max for the 12-MRT is estimated from the following formula, where *d*_12_ is the distance covered in meters:^15^ 

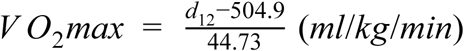

### Heart Rate Calibration Study Procedures and Measures

All study procedures were approved by the UCSD Institutional Review Board (approval number 181820). All participants provided written informed consent and attended one in-person study visit at EPARC.

A convenience sample of 120 adults, 18-65 years old, of six various skin types were asked to participate in this study. We aimed to recruit an equal ratio of male and female participants, as well as an equal number of each skin type as determined by the Fitzpatrick scale. Participants were included if they were 1) able to consent and participate in the study using English; and 2) between 18 and 65 years of age. Participants were excluded if they had 1) peripheral neuropathy; or 2) tattoos or scarring at the measurement site (index finger and/or wrist). Potential participants were contacted by trained EPARC staff via email or telephone, and they were asked to complete screening to ascertain their eligibility.

To establish the Fitzpatrick skin type of the cohort during recruitment, participants were asked to self-assess their Fitzpatrick skin type based on visual comparison to images of well-known celebrities with diverse pigmentation levels. As self-assessment of skin type can have variable accuracy,^31,32^ spectrocolorimetry was also used as an objective standard.^33^ Spectrocolorimetry measurements were performed on the underside of the jaw using a Pantone RM200QC. To calculate pigmentation in the Individual Typology Angle (ITA) color space, the L* parameter and the b* parameters from the spectrocolorimetry measurements were used according to the formula: °ITA = [arctan((L*−50)/b*)] × 180/3.14159. Using this formula, skin color types can be classified into six groups, ranging from very light to dark skin: very light > 55° > light > 41° > intermediate > 28° > tan > 10° > brown > 30° > dark.^33^

Upon completion of the telephone screening, potential participants were scheduled to attend the first testing session at UCSD. Participants were asked to provide their age, sex at birth, ethnicity, and race. All participants were fitted with a chest-worn heart rate monitor (Polar, Finland) that was used for real-time monitoring by trained EPARC staff throughout testing. Heart rate was also monitored using a finger-based pulse oximeter (Nonin Medical, Inc. Plymouth, MN). The finger-based pulse oximeter was attached to the participants’ index finger and the time was synced between the computer and the device. Trained research staff visually confirmed that the photoplethysmograph was reading accurately before starting measurement on smartphone devices.

Participants were then given the first of eight smart phones: Huawei Mate SE, LG Stylo 4, Moto G6 Play, Samsung Galaxy J7, Samsung Galaxy S9+, iPhone8+, iPhoneSE, iPhoneXS. They were instructed by trained research staff to stand still, and gently cover the camera and flash on the back of the smartphone with their fingertip as their heart rate was captured by our preloaded smartphone application. The time on the Polar application was recorded at the time the measurement began on the smartphone application. Measurement with each smartphone lasted 60 seconds in duration. Processed data from the finger-based pulse oximeter was parsed and transformed with custom scripts to generate continuous PPG data in a format suitable for comparison with the heart rates from the phones.

## Statistical analysis

Demographic data was described using univariate summary statistics (e.g. proportions, mean, and standard deviation). Test validity for heart rate estimates and VO_2_max was visualized using Bland-Altman plots^17^ and compared using Lin’s concordance index^34^. Heart rates errors were also compared using percent error. Analysis was carried out in both R and Python.

### iOS and Android Heart Snapshot Software Modules

Code for the Heart Snapshot modules and sample Android (https://github.com/Sage-Bionetworks/CardiorespiratoryFitness-Android) and iOS (https://github.com/Sage-Bionetworks/CardiorespiratoryFitness-iOS) applications are available under an open source license.

